# RASflow: An RNA-Seq Analysis Workflow with Snakemake

**DOI:** 10.1101/839191

**Authors:** Xiaokang Zhang, Inge Jonassen

## Abstract

**Background:** With the cost of DNA sequencing decreasing, increasing amounts of RNA-Seq data are being generated giving novel insight into gene expression and regulation. Prior to analysis of gene expression, the RNA-Seq data has to be processed through a number of steps resulting in a quantification of expression of each gene / transcript in each of the analyzed samples. A number of workflows are available to help researchers perform these steps on their own data, or on public data to take advantage of novel software or reference data in data re-analysis. However, many of the existing workflows are limited to specific types of studies. We therefore aimed to develop a maximally general workflow, applicable to a wide range of data and analysis approaches and at the same time support research on both model and non-model organisms. Furthermore, we aimed to make the workflow usable also for users with limited programming skills.

**Results:** Utilizing the workflow management system Snakemake and the package management system Conda, we have developed a modular, flexible and user-friendly RNA-Seq analysis pipeline: RNA-Seq Analysis Snakemake Workflow (RASflow). Utilizing Snakemake and Conda alleviates challenges with library dependencies and version conflicts and also supports reproducibility. To be applicable for a wide variety of applications, RASflow supports mapping of reads to both genomic and transcriptomic assemblies. RASflow has a broad range of potential users: it can be applied by researchers interested in any organism and since it requires no programming skills, it can be used by researchers with different backgrounds. RASflow is an open source tool and source code as well as documentation, tutorials and example data sets can be found on GitHub: https://github.com/zhxiaokang/RASflow

**Conclusions:** RASflow is a simple and reliable RNA-Seq analysis workflow which is a full pack of RNA-Seq analysis.

## Background

RNA sequencing (RNA-Seq) was introduced more than ten years ago and has become one of the most important tools to map and identify genes and understand their regulation and roles across species [1, 2]. A large number of studies have been performed using RNA-Seq and resulted in gene expression data sets available in databases such as GEO [3] and ArrayExpress [4]. Underlying reads are typically deposited to the Sequence Read Archive [5], currently containing reads for more than 1,5 million samples (https://www.ncbi.nlm.nih.gov/sra/?term=RNA-Seq). One of the most popular applications of RNA-Seq is for Differential Expression Analysis (DEA) where one identifies genes that are expressed at different levels between two classes of samples (e.g., healthy, disease) [6].

When RNA-Seq is used in a DEA project, the reads coming out of the sequencer need to be taken through several steps of processing and analysis. Often, the steps are organized into a workflow that can be executed in a fully or partially automated fashion. The steps include: quality control (QC) and trimming, mapping of reads to a reference genome (or transcriptome), quantification on gene (or transcript) level, statistical analysis of expression statistics to identify genes (or transcripts) showing differential expression between two pre-defined sets of samples, along with associated p-values or False Discovery Rate (FDR) values. Mapping reads to the genome is the most computationally intensive and time-consuming step. An alternative approach performing pseudo alignment to a transcriptome, has gained more popularity recently, due to its high speed and high accuracy [7, 8, 9]. It has been shown that lightweight pseudo alignment improves gene expression estimation and reduces computational consumption compared with the standard alignment/counting methods [10]. But if the purpose of analysis is to call variants instead of DEA, then it is still better to map reads to genome to avoid reads mapped to spurious locations on the transcriptome [11]. Considering this, a workflow should provide both quantification strategies to satisfy users with different research interests.

There is a large number of RNA-Seq analysis workflows and many have been published and made available for the user community. We reviewed 7 workflows published in the past three years. We find that none of these are as comprehensive and flexible as would be desirable to cover a large range of user groups and needs. So a more complete and also easy-to-use workflow is still needed.

In this article we present RNA-Seq Analysis Snakemake Workflow (RASflow) that is able to cover many needs for a wide range of different users. RASflow can be applied to any organism and can map reads to either a genome or transcriptome, allowing the user to refer to public databases such as ENSEMBL [12] or to supply their own genomes or transcriptomes [13, 14]. The latter can for example be useful for projects on non-model species for which there is no public high-quality reference genome/transcriptome. RASflow is scalable: it can be run on either supercomputers with many cores (which enable parallel computing) or on a personal laptop with limited computing resources; it can process data from up to hundreds of samples and still consume very little storage space because it temporarily copies or downloads the FASTQ file(s) of one sample (one file for single end and two files for pair end) to the working directory at the time, and it stores only the necessary intermediate and final outputs. Using Conda, the whole workflow with all dependencies (version already specified) can be installed simply with one single command in a virtual environment. This ensures a quick and smooth installation. Using Snakemake [15], the whole analysis is completely reproducible and highly user-friendly also for users with limited programming skills. In the DEA step, RASflow supports use of paired tests - that can help to strengthen the statistical power and bring out expression differences related to the phenomenon under study [16].

## Implementation

The workflow of RASflow is shown as Fig. 1. It starts with QC of the raw FASTQ files using FastQC (https://www.bioinformatics.babraham.ac.uk/projects/fastqc/). The QC report will be presented to the user and the user will be asked whether reads should be trimmed. If yes, the reads will go through trimming using Trim Galore (https://www.bioinformatics.babraham.ac.uk/projects/trim_galore/). After trimming, another QC report will be generated.

**Figure 1.**
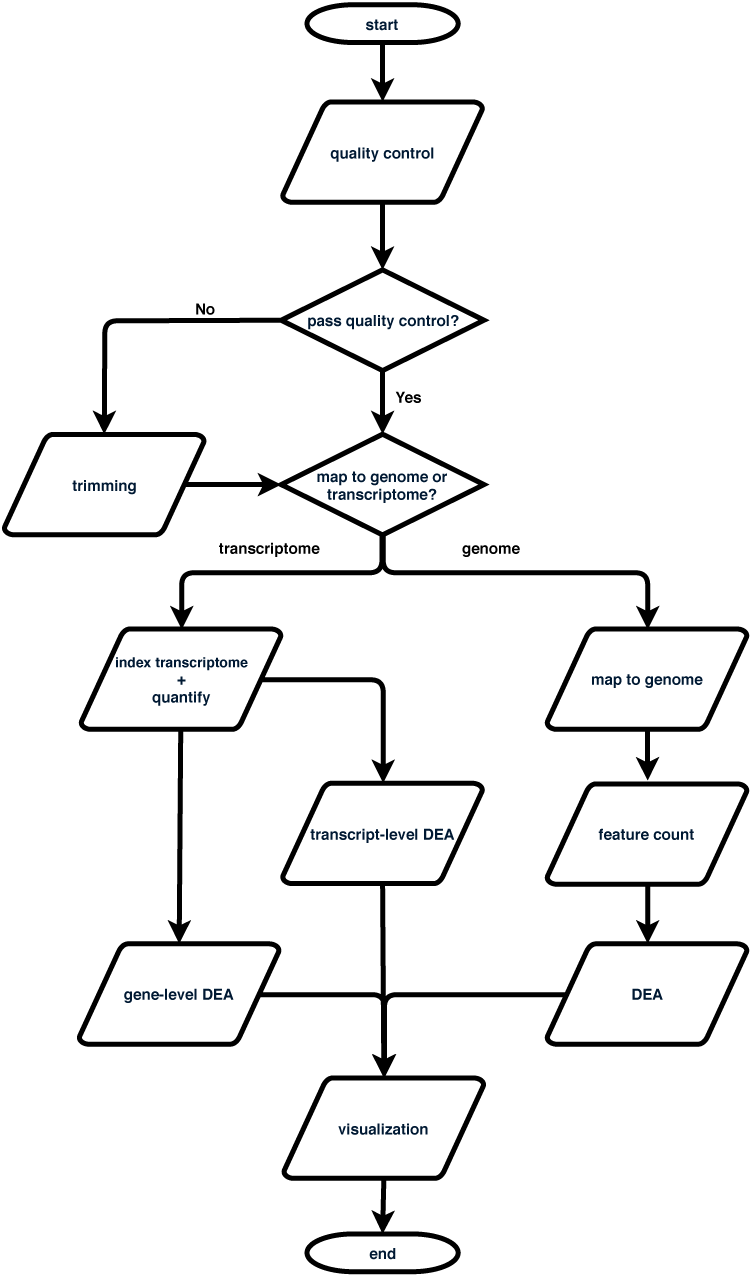
Overview of the steps performed by RNA-Seq Analysis Snakemake Workflow (RASflow).

When the user is happy with the quality of the data, the workflow proceeds to the next step: quantification of read abundance / expression level for transcripts or genes. The user needs to decide whether to map the reads to transcriptome or genome based on their goal of the analysis. If it is for DEA, it is suggested to map the reads to transcriptome using pseudo alignment with Salmon [9]. Pseudo alignment is less computing consuming and makes it possible to do DEA not only on gene level but also on transcript level which sheds light on the transcript dynamics. A quantification table of the transcripts will be generated from this step. Alternatively, the user can choose for the reads to be mapped to a genome which is more robust and is also a requirement for some types of downstream analysis, such as variant calling. The aligner used in RASflow is HISAT2 [17] which has relatively modest memory requirements (∼4.3GB for the human genome) compared with for example the STAR aligner (requiring ∼32GB for the human genome) [18]. The alignment is followed by a quality evaluation done by Qualimap2 [19] and feature count done by featureCounts [20]. To be noted, after most of the steps, a summary report will be generated using MultiQC [21].

After the reads have been mapped to the chosen transcriptome / genome, RASflow proceeds to perform a DEA analysis using the edgeR tool [22, 23]. This tool is capable of doing paired test for experiments designed in this way. The user can easily specify the statistical test mode in the configuration file of RASflow. If the reads were mapped to transcriptome, DEA will be done on both transcript- and gene-level. In any case, the outputs of DEA include three types of tables: normalized quantification tables, important statistics (such as Log Fold Change, FDR) for the whole gene or transcript list, and the list of significantly differentially expressed genes or transcripts (with default threshold of *FDR* < 0.05). For the normalized quantification tables, if the reads were mapped to genome, then the raw count is normalized based on Trimmed Mean of M values (TMM) [24]; if the reads were mapped to transcriptome, then the values in the normalized quantification tables are actually estimated Transcripts Per Million (TPM) from Salmon but scaled up to library size using “tximport” [25]. The results of DEA will also be visualized with a volcano plot which gives a visual identification of genes with high fold change that are also statistically significant, and a heatmap which not only visualizes the expression pattern of selected differentially expressed genes but also uses that to cluster the samples, so that the user can get an idea of how well the groups are distinguished.

## Results

To show the users how RASflow works and to familiarize them with RASflow, we provide some small example data sets. They are generated as subsets of the original real data [26]. The figures in this section were generated by RASflow using the example data as input.

### Quality control

FastQC checks the quality of the sequencing reads and gives one report for each FASTQ file, and MultiQC summarizes all the reports and merge them into one report, as shown in Fig. 2a and Fig. 2b.

**Figure 2.**
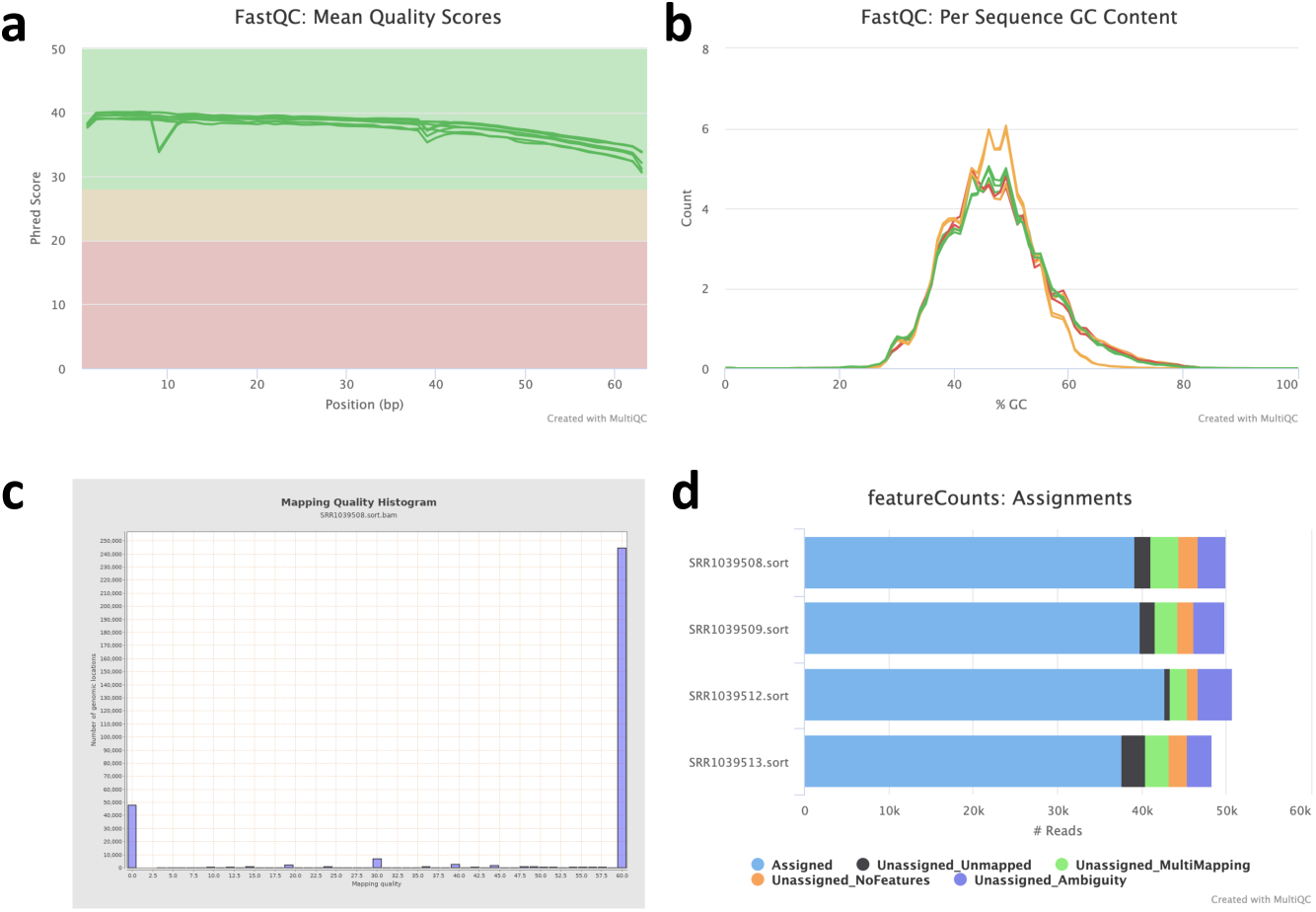
Quality control of raw reads and alignment. **a** The mean quality value across each base position in the read. **b** The average GC content of reads. Normal random library typically have a roughly normal distribution of GC content. **c** Histagram of mapping quality. **d** A brief mapping summary.

After the alignment to genome, the intermediate output, bam files will be provided to Qualimap2 to evaluate the alignment quality. Fig. 2c shows one of the reports from Qualimap2.

MultiQC also generates a report using the output of featureCounts which gives some statistics of the mapping ratios (Fig. 2d).

### Differential expression analysis

RASflow can satisfy both single- and paired-test with edgeR. Choose the proper one based on the experiment design.

Besides quantification tables and important statistics of the gene or transcript, RASflow also generates some visualization such as volcano plot (Fig. 3a) and heatmap (Fig. 3b).

**Figure 3.**
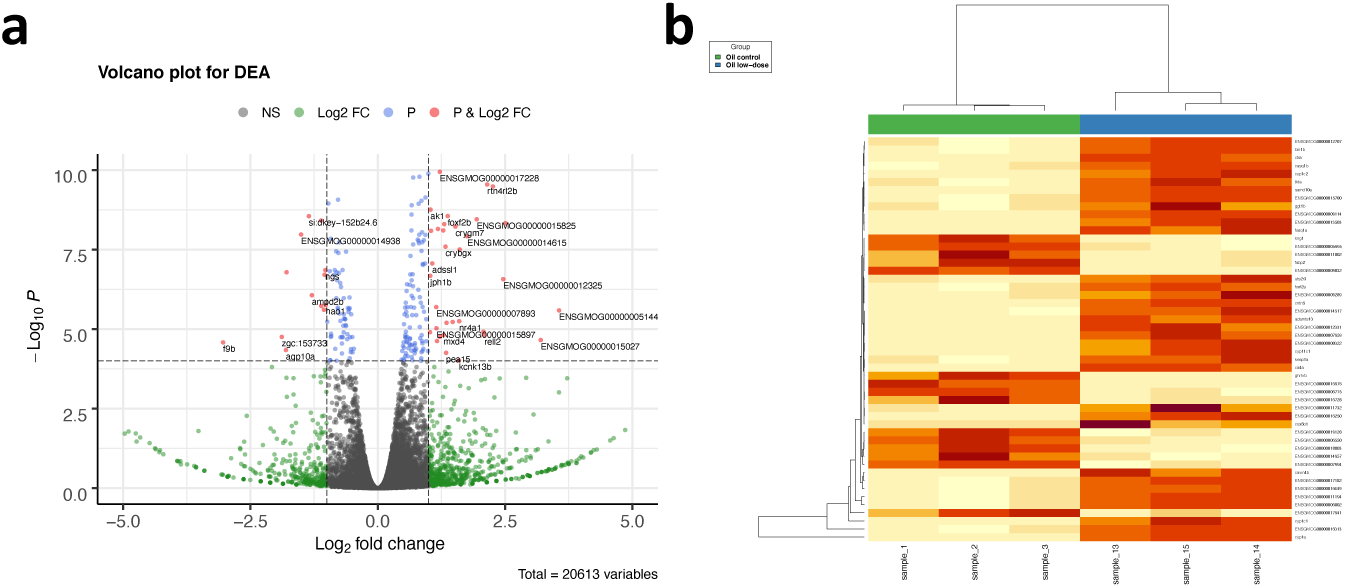
Visualization of DEA results. **a** Volcano plot with labeled genes who pass the thresholds of both Fold Change and P-value. **b** Hierarchical clustering heatmap with samples along the x-axis and differentially expressed genes along the y-axis.

## Discussion

### Snakemake as framework

Snakemake is a scalable workflow engine which helps to manage workflow steps in an easy way. It divides the whole workflow into rules with each rule accomplishing one step. The input of a rule is the output from the rule of the previous step, making the data flow easy to track. Thanks to this logic, the whole workflow becomes highly modular, so users can easily expand the workflow or replace part of it, no matter how complicated it is.

RASflow organizes the rules into several files so that the rules in one file work together to realize one function (one big step in the whole workflow). Each file has one corresponding configuration file where the users specify the important parameters, such as the directory of the read files.

### Transciptome and genome as reference

RASflow allows users to supply their own genomic or transcriptomic reference. This enables users to study expression in species where no public reference is available or the users have alternative references that they wish to utilize. It should be noted that if one aims for transcript-level analysis, a transcriptome should be used as reference.

But some analysis other than DEA requires the reads to be mapped to genome and gene-level differential expression analysis is more robust and experimentally actionable, so RASflow still provides the traditional workflow of genome alignment and differential expression analysis based on gene counts.

### Comparison to other tools

We compared RASflow to other existing workflows as shown in Table 1. As we can see from the table, some workflows do not include all the steps, either missing the QC steps [27, 28], or the last steps, quantification and DEA [29]. Some of the workflows are limited to specific organisms typically human or mouse and in some cases other model organisms [30, 27, 31, 32]. Some of them have functionality only for mapping reads to a reference genome and do not support the use of a transcriptome reference [27, 31, 32, 33]. Most of the workflows have high demands: some of them require a high-performance computer with large internal memory [31, 29, 32, 33]; some of them need complex installation processes depending on the user to install other tools or libraries separately, which easily leads to version conflicts [28, 33]; some workflows require users to be able to program which makes them unusable for some biologists [28, 33]. Those issues limit the wide-range use of the workflows. Those are the motivation of the development of RASflow which overcomes those drawbacks.

**Table 1.**
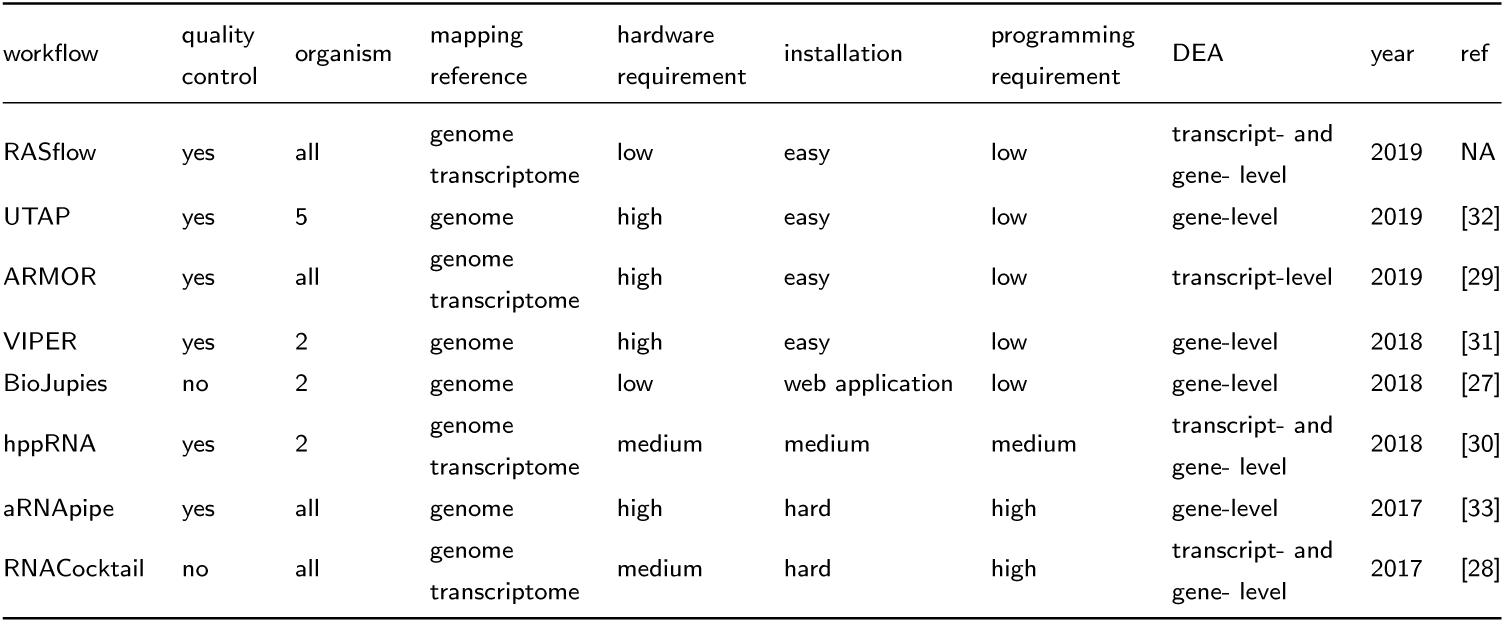
Comparison of RASflow with the other workflows published between 2017 and 2019.

### Extension of RASflow

Thanks to the high modularity of RASflow, it is very easy to change the tools applied in RASflow. For example, after mapping the reads to the genome, the user can choose to use either featureCounts or htseq-count [34] to count the features by simply specifying this option in the configuration file. Advanced users can also replace any tool with their favorite one.

RASflow can also be extended to realize other functions, such as Single Nucleotide Polymorphism (SNP) detection, pathway analysis, and so on.

## Conclusions

RASflow is a light-weight and easy-to-manage RNA-Seq analysis workflow. It includes the complete workflow for RNA-Seq analysis, starting with QC of the raw FASTQ files, going through trimming (alternative, depending on the QC report), alignment and feature count (if the reads are mapped to genome), pseudo alignment (if transcriptome is used as mapping reference), gene- or transcript-level DEA, visualization of the output from DEA.

RASflow is designed in such a way that it can be applied by a wide range of users. It requires little programming skills from the users and a well written tutorial to help users go through the whole workflow makes it very easy to set up and run RASflow from scratch. RASflow has a very low hardware requirement so that it can almost be run on any personal computer. It can also be scaled up to make full use of the computing power if it is run on a supercomputer or cluster. RASflow can be applied to any organism and the users can also choose to map the reads to genome or transcriptome. Especially when the organism is not a model organism and no public genome or transcriptome is available or the users want to use an alternative reference, they can use RASflow to map the reads to their own reference.

RASflow is built on the basis of Conda and Snakemake, which makes the installation and management very easy. All the required tools are available on Anaconda cloud (https://anaconda.org/) and are wrapped in a virtual environment managed by Conda, so RASflow is independent of the system and package / library version conflicts are therefore avoided. The whole workflow is defined by rules managed by Snakemake, which makes it highly modular. This means that the advanced users can easily extract part of the workflow or expand it based on their own research needs, and replace the tools used in RASflow with other tools to explore new pipelines for analyzing RNA-Seq data.

## Availability and requirements

Project name: RASflow.

Project home page: https://github.com/zhxiaokang/RASflow

Operating system(s): Linux and MacOS.

Programming language: Python, R, Shell

Other requirements: Conda

License: MIT License

Any restrictions to use by non-academics: N/A.

## List of abbreviations

RASflow: RNA-Seq Analysis Snakemake Workflow;
DEA: Differential Expression Analysis;
QC: Quality Control;
TMM: Trimmed Mean of M values;
TPM: Transcript Per Million;
FDR: False Discovery Rate;
SNP: Single nucleotide polymorphism.

## Ethics approval and consent to participate

Not applicable.

## Consent for publication

Not applicable.

## Availability of data and materials

All the datasets and source codes are available on GitHub.

## Competing interests

The authors declare that they have no competing interests.

## Funding

The dCod 1.0 project is funded under the Digital Life Norway initiative of the BIOTEK 2021 program of the Research Council of Norway (project no. 248840).

## Author’s contributions

XZ designed and developed RASflow and wrote the tutorial. XZ wrote the initial draft of the manuscript. IJ supervised the work and finalized the manuscript.

## Acknowledgements

We would like to thank the colleagues in dCod 1.0 project for which the workflow was initially designed. The feedback from the biologists in the project greatly helped the improvement of the workflow. We would also like to thank the research school NORBIS where XZ took the training on analysis of next generation sequencing.

## Notes

https://github.com/zhxiaokang/RASflow

